# Tuning of recombinant protein expression in Escherichia coli by manipulating transcription, translation initiation rates and incorporation of non-canonical amino acids

**DOI:** 10.1101/100982

**Authors:** Orr Schlesinger, Yonatan Chemla, Mathias Heltberg, Eden Ozer, Ryan Marshall, Vincent Noireaux, Mogens Høgh Jensen, Lital Alfonta

## Abstract

Protein synthesis in cells has been thoroughly investigated and characterized over the past 60 years. However, some fundamental issues remain unresolved, including the reasons for genetic code redundancy and codon bias. In this study, we changed the kinetics of the E. coli transcription and translation processes by mutating the promoter and ribosome binding domains and by using genetic code expansion. The results expose a counterintuitive phenomenon, whereby an increase in the initiation rates of transcription and translation lead to a decrease in protein expression. This effect can be rescued by introducing slow translating codons into the beginning of the gene, by shortening gene length or by reducing initiation rates. Based on the results, we developed a biophysical model, which suggests that the density of co-transcriptional translation plays a role in bacterial protein synthesis. These findings indicate how cells use codon bias to tune translation speed and protein synthesis.

---

Protein synthesis, one of the most important and complex functions of living cells, is controlled by several mechanisms. Every stage in the process, from DNA transcription to protein folding dynamics, is tightly regulated to ensure that proteins are produced in required amounts, at the correct times and with minimal waste of energy and resources^1^. In bacteria, the transcription of DNA to mRNA and the subsequent translation into a polypeptide chain are coupled in time and space ^2,3^. The two processes occur simultaneously, which creates a high molecular density area populated with all the components required for protein synthesis. For the dynamics of transcription and translation, such molecular crowding in the cytoplasm plays an important role by stabilizing protein-protein interactions and by controlling the diffusion rates of the components involved in protein synthesis^
4,5^. The molecular densities of RNA polymerases on DNA and of ribosomes on mRNA are known to depend on the transcription and translation initiation rates, which, in turn, are determined by the strengths of the promoter and of the ribosome binding site (RBS). For example, it was shown that the use of a strong RBS with a high initiation rate to overexpress proteins can lead to ribosome collisions and queuing along individual mRNA strands. These queues can generate interference between adjacent translating ribosomes, significantly lowering the yields and efficiency of protein expression^6,7
^. The nature of possible interactions that may occur between ribosomes on adjacent mRNA strands, however, is not clear.

The kinetics of translation also depend on the encoded gene’s codon bias, which is manifested by its effects on the elongation rate of the growing polypeptide chain^8,9^. Exploited across species to control translation rates and the ribosome queues along mRNA strands, codon bias is used to optimize protein synthesis and folding. Depending on the elongation rates they dictate, codons can be divided into different rate classes. Slower codons are found to be more favorably encoded for in the first 30-50 codons of the mRNA, thus resulting in ribosome crowding near the translation initiation site. Downstream codons, however, are found to be optimized for fast elongation rates ^10–12^. These findings give rise to several questions: Why is translation that occurs close to the translation initiation site slow? Is this slow translation rate related to the density of the molecular environment in the vicinity of the co-transcriptional-translation event?

Although normally used for applicative purposes,^13^ genetic code expansion through stop codon suppression constitutes an effective, basic research tool to shed light on these questions. One approach of genetic code expansion, the incorporation of non-canonical amino acids (ncAAs) into proteins, typically exploits the UAG nonsense (stop) codon, essentially transforming it into a sense codon that encodes for the incorporation of a ncAA. This recoding is facilitated by introducing into a host organism an orthogonal translation system (OTS) that comprises an orthogonal *archaeal* o-tRNA with an anticodon corresponding to the UAG stop codon and an orthogonal amino-acyl-tRNA synthetase (o-aaRS) that selectively recognizes the ncAA of choice and aminoacylates its cognate tRNA_CUA_^14^. The affinity of the o-tRNA to the tertiary complex of the ribosome A-site during translation, significantly smaller than that of the native bacterial tRNA ^15,16^, can be exploited to alter ribosomal traffic on the mRNA by decreasing the speed of translation along the mRNA. This approach can only succeed, however, when the OTS and the native release factor (i.e., RF1) are not in direct competition for the UAG codon. That competition can be eliminated by recoding all TAG stop codons in the bacterial genome to TAA and by knocking out the RF1 gene ^17^.

Here we use the OR2-OR1-pr-UTR1 (P70a-UTR1) expression system, based on a modified lambda PR promoter and the T7 bacteriophage RBS ^18^, to perform genetic code expansion. This system has the highest transcription and translation initiation rates reported for an *E. coli* element, and so far, it has been used exclusively *in-vitro*. Its high initiation rates promote large and unusual ribosome crowding along the transcribing mRNA. We therefore hypothesized that in the crowded environment of a polysome, a growing polypeptide chain may interact with neighboring translational components inside the polysome in a manner that can significantly retard the process. Indeed, it was previously shown that the nascent polypeptide can regulate the translation process in the ribosome by interacting with the polypeptide exit tunnel in the ribosome^19^. Such interaction may cause ribosome stalling ^20^, translation arrest ^21^ and even accelerated mRNA degradation ^22^. We exploited both the incorporation of ncAA using UAG stop codon suppression, synonymous mutations in the gene and the modular tuning of the P70a-UTR1 expression system to model and control ribosomal traffic, thus optimizing recombinant protein expression.

## Results

### WT GFP exhibits smaller expression levels compared to GFP with ncAA incorporated at position 35

Compared to its *in vitro* expression, the *in vivo* expression of the GFP (WT GFP stands for protein without incorporated ncAAs) using the strongest *E. coli* promoter so far reported (P70a-UTR1) ^18^ was unexpectedly weak (Figs. 1A, 1B lane b). This outcome was observed not only when using a genomically recoded *E. coli* strain (C321Δ*prf1*) (GRO)^17^, but also with several other *E. coli* strains (i.e., BL21(DE3) & DH5α). However, expression in the GRO strain of the same protein, in which a tyrosine residue at position 35 had been replaced with a nonsense stop codon (UAG), led to large and unexpected quantities of mutant GFP with an unnatural amino acid incorporated into position 35 (Figs. 1A, 1B lane d). Correct ncAA incorporation was verified by mass spectrometry (LC-ESI-MS) as well as by MS/MS analysis of peptide fragments (Fig. S1).

To understand these initial observations, we first ruled out the possibility that inclusion bodies or secondary mRNA structures were the source of the divergence between the WT GFP and 35TAG GFP quantities. Cryo-electron microscopy (Cryo-EM) imaging of GFP revealed neither inclusion bodies nor any marked difference in bacterial shape compared to Cryo-EM images of bacteria without the GFP expression plasmid (Figs. S2A, S2B). Moreover, there was no difference in the mRNA structure encoding for the WT GFP and the mutant GFP (Figs. S2C, S2D). We used two different vector plasmids to express the mutant Y35TAG GFP: one encoding for the mutated protein and one encoding the Pyrrolysine Orthogonal Translation System (Pyl-OTS), which is the machinery for the ncAA incorporation. To exclude the possibility that the pEVOL Pyl-OTS plasmid contributed to the unusual overexpression of the mutated reporter protein, we demonstrated that pEVOL Pyl-OTS has no particular effect on the expression of WT GFP (Fig. S2E). Taken together, these observations motivated our search for a more fundamental explanation related to the coupling of bacterial transcription and translation kinetics.

### Density induced translation arrest model predictions corresponds to counterintuitive protein expression patterns

Herein, we propose a model to predict protein and mRNA levels that is based on a set of biochemical parameters combined with several assumptions. Model parameters: An increase in the RNA polymerase (RNAP) initiation rate (i.e., promoter ‘strength’) leads to a decrease in the average distance between transcribing RNAP and vice versa ^23^. The deterministic average distance between RNAPs, <D>, is governed by equation (1) (The analytic equation and its solutions are presented in Figs. S3A, S3B):

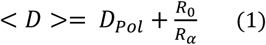

where R_α_ is the RNAP initiation rate and R_0_ is the RNAP elongation rate anywhere on the gene, while D_Pol_ is the size of the RNAP, which defines the minimum distance between polymerases. The use of the Gillespie stochastic algorithm imposed a distribution of RNAP velocities around the simplified elongation rate: R_0._ This creates a stochastic distribution of the distances between RNAPs and even creates queues of adjacent RNAPs. As the average distance between RNAPs decreases, the density of mRNAs being synthesized along the DNA strand increases and the average distance between adjacent mRNAs decreases. We named this promoter dependent mRNA density along DNA as ‘**transcriptional density**’ (Fig. S3E).

The initiation rate of translation is directly proportional to ribosome affinity to the ribosome binding site (RBS). As the ribosomal translation initiation rate increases, the average distance between the ribosomes translating the same mRNA template becomes shorter. The average distance, <d>, is governed by ribosome size d_rib_, the initiation rate r_α_ and the elongation rate for each codon ‘i’ given by r_i_’. Considering that the time for each step is given by 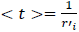 the average distance (in number of codons) can be simplified and expressed by equation (2) (The analytic equation and its solutions are presented in Figs. S3C, S3D):

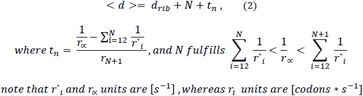

Another factor to include in the model is the different elongation rates of each codon in the mRNA sequence ^24–26^. Based on a previous model developed by Mitarai et al., the entire set of bacterial codons were divided into three groups based on translation rate – fast (A), medium (B) and slow (C) – which correspond to elongation rates of 35, 8 and 4.5 codons per second, respectively^25^ (Fig. S3F). To these canonical codons we added the new non-canonical UAG codon (only translated in the GRO strain by the o-tRNA), which significantly lowered the codon elongation rate. The UAG codon was assigned a new translation rate category, group (D), which had a significantly lower elongation rate of 0.04 codons per minute. The rate was estimated from in vitro experiments ^15^ and even though this value has some uncertainty to it, it is at present our best estimate. Moreover, the model based simulated results are quite robust to large perturbations around this estimate. For example, the main observation being that by using an early TAG mutation, significantly more protein is being produced compared to WT GFP. These yields are still achieved for values of a UAG rate ranging between 0.01 codons/s - 0.2 codons/s. Like the case of the RNAP stochastic velocity, the ribosome also moves in a stochastic-probabilistic manner. This means that in addition to the 4 rate groups the actual ribosome velocity is governed by rate distributions for each codon around the group mean. Finally, we included ‘**translational density**’, defined as the density of ribosomes along an mRNA. The length of the growing nascent polypeptide is directly proportional to the position of the ribosome along the mRNA relative to the translation initiation site.

In bacteria, transcription and translation are coupled, i.e., as soon as the RBS on the transcribing mRNA emerges from the RNAP, the ribosome binds the RBS and begins translation ^
2
^. The close proximity of the two processes in time and space means that there may be interactions between them. Accordingly, we hypothesized that highly crowded conditions will promote random interactions between nascent polypeptides and ribosomes, thus inducing translation arrest in a process that we termed ‘Density Induced Translation Arrest’ (DITA). We propose that in cases in which the promoter and RBS initiation rates are large enough to create regions with high molecular density and in which the nascent polypeptide is long enough, the probability for DITA events increases. In the case of a DITA event, all the ribosomes upstream of the arrested ribosome stall, promoting translation termination and thus reducing the number of full-length proteins produced from crowded mRNA strands (Fig. 1C).

Next, we characterized our system’s model parameters (described in detail in the methods section and listed in Table S2). The GFP gene was mapped and the codons were assigned to one of the four codon rate groups (A-D). The ribosome elongation rate is governed by each codon during translation. The average RNAP transcription rate was assumed to be constant ^2,27^. Lastly, the length of the growing nascent polypeptide could not be determined a priori since its folding dynamics and interactions with the ribosome are unknown to us. For this reason, we chose the simplest possible approach and we added an empirical constant of proportionality, λ, which governs the length of the polypeptide protruding from its parent ribosome in the model (Fig. S3H). This approach allowed us to predict, for a given gene, which transcription-translation instances will generate a full-length protein and, as a result, the protein production rate.

The results of the Gillespie algorithm simulation agreed with the experimental results for both WT and Y35TAG mutant GFP (Fig. 1A). The model suggests that WT GFP expression levels are negligible because of the high probability for DITA occurrences when a strong promoter and RBS, such as P70a and UTR1, respectively, are used. In the case of the Y35TAG GFP mutant, the model suggests that the small-translation-rate UAG codon (group D) inserted in this position serves as a 'traffic light' that reduces ribosomal density downstream. Taken together, the reduction in translational density downstream of the UAG codon and the low probability of a DITA event result in high yields of expressed protein.

**Figure 1.**
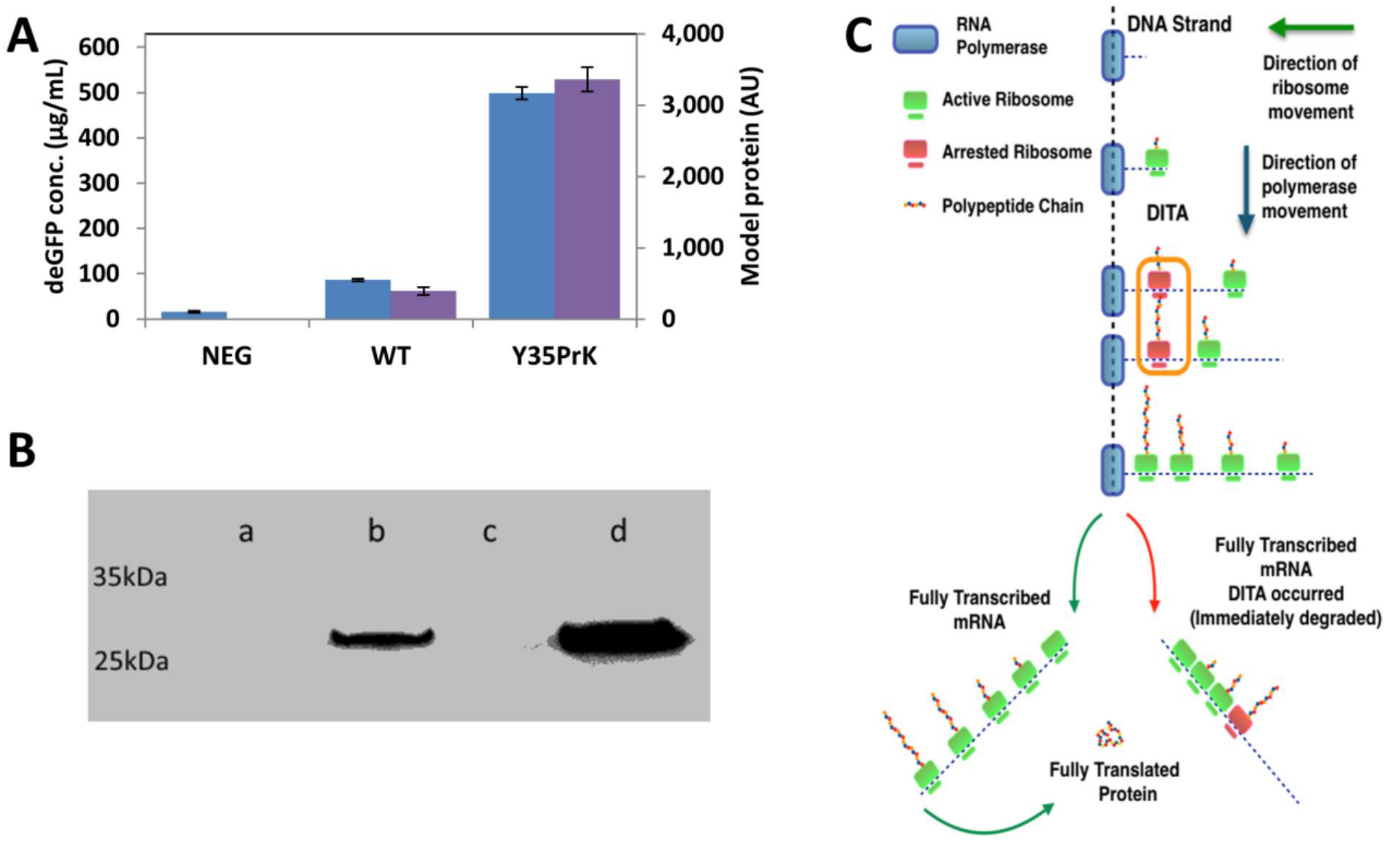
GFP expression using the P70a-UTR1 system. (A) Comparison of the experimental results(blue bars) with the modeled protein quantities (purple bars). (B) Western blot analysis using anti-GFP antibody of GFP expression in C321.ΔA.exp: w/o plasmids(a), pBEST-p70a-UTR1-GFP WT (b) and Y35PrK mutant in the absence and presence of PrK in the growth medium (lanes c and d, respectively). (C) Schematic presentation of the hypothesized DITA phenomenon.

### Early reassigned Amber stop codon rescues protein expression as predicted by the model

In the proposed model, we suggest that a suppressed UAG stop codon functions as a traffic light, thus its position along the mRNA is of importance. Due to its substantially slower ncAA incorporation kinetics compared to those of codons encoding for canonic amino acids, a queue of ribosomes will grow behind the reassigned stop codon. The transient stalling generated by an early UAG codon significantly reduces ribosome occupancy downstream, thereby reducing the chance of a DITA event (Fig. 2A). As the translation process continues, the chance that the elongating polypeptide chain will have a DITA grows. Indeed, both our experimental results and our simulations indicated that the earlier the stop codon is introduced, the lower the chance of a DITA event. For cases in which both promoter and RBS are strong, our hypothesis predicts that the closer a UAG codon is positioned to the C-terminal, the smaller will be the protein yields in a manner similar to what is observed for the WT GFP. Indeed, the choice of a late D193TAG site in the simulation resulted in high DITA levels and small protein yields compared to those in the Y35TAG GFP mutant and protein yields equal to that of the WT protein. To test our prediction, we mutated position D193TAG in GFP. The experimental results coincided with those of the simulation, i.e., low protein levels (Fig. 2B). Note that D193TAG GFP is a permissive mutation site, as compared *in-vitro* to the WT and Y35PrK mutant expression (Fig. S4A). The relationship between the position of the UAG codon and the protein expression levels was simulated (Fig. S4B) revealing that only the first 37 codons enable rescue of protein levels. This result is in agreement with our experimental results and with earlier reports by Tuller et. al of an early slow translating “ramp” region close to the translation initiation region^
9,10
^.

**Figure 2.**
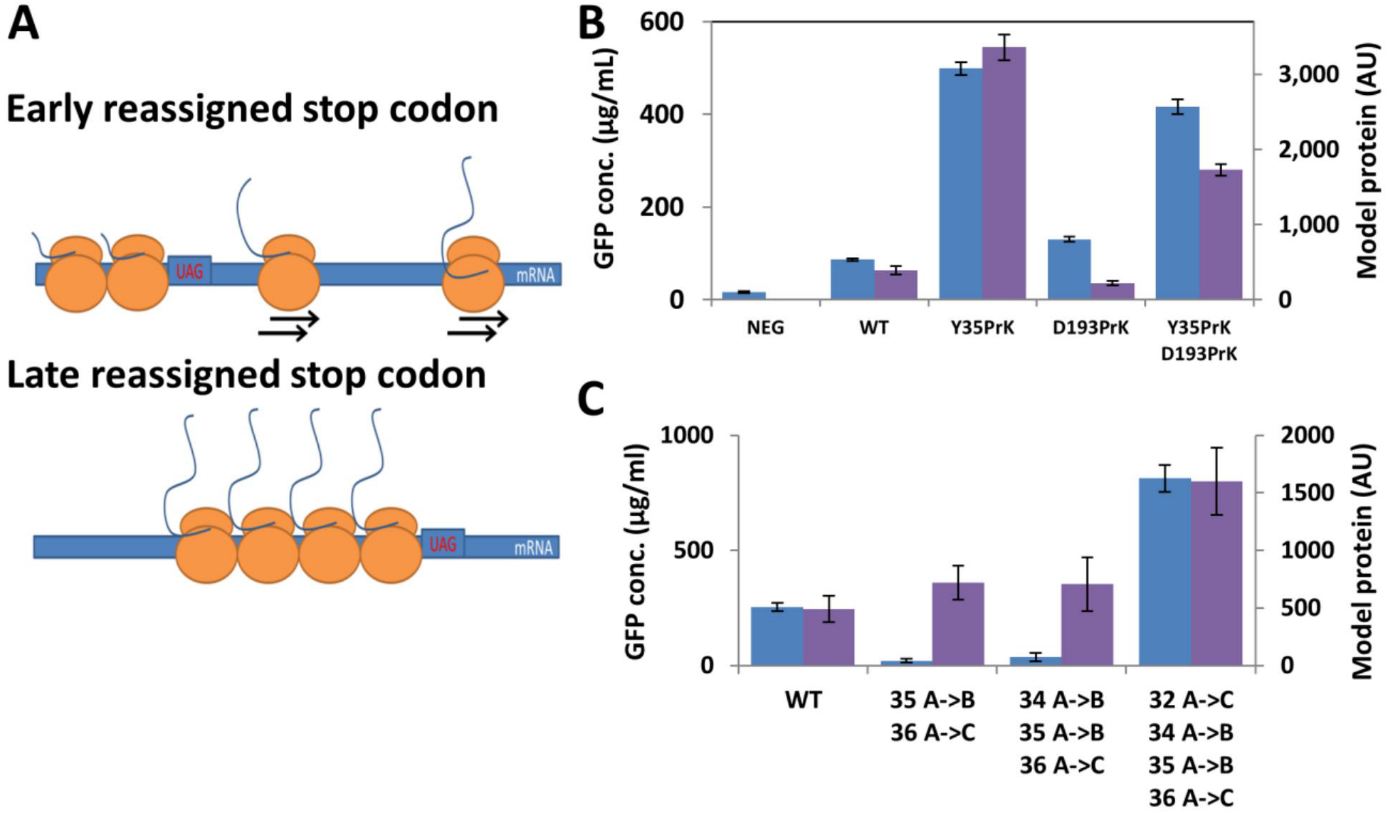
The ‘rescue effect’ caused by ncAA incorporation and the associated attenuation in transcription rate is position dependent as well as “slow translating” codon dependent. (A) Schematic presentation of the ribosome queue caused by UAG codons at various positions along the mRNA. In early introduced UAG codons, the nascent polypeptide is relatively short and the ribosome traffic downstream of the codon is low, reducing the chance of peptide interactions. However, in later UAG codons, the nascent peptide is longer and has more time to interact with neighboring peptides in the polysome due to the queue caused by the slow rate of the UAG codon. (B) Comparison of the experimental results (blue bars) with the modeled protein amounts (purple bars). (C) WT GFP expression upon introduction of synonymous mutations around position 35. Experimental results (blue bars) and modeled protein quantities (purple bars) show a significant increase in expression after the introduction of the 4^th^ synonymous mutation.

Next, we tested the influence of adding an early UAG codon to a mutant that already contains a late mutation (Y35TAG+D193TAG). The model predicted that the early mutation would decrease the translational density around the later UAG stop codon, thus reducing the probability of DITA and conferring a rescue mechanism on protein levels. The expression levels of the double mutant Y35TAG+D193TAG GFP, its protein expression kinetics and the final yields with the different mutants predicted *in-silico* and tested *in-vivo* showed high correlation and a clear rescue effect on protein expression (Fig. 2B).

Observing these results, we wanted to test whether the rescue effect could be achieved with synonymous (silent) sense codon mutations. Hence, we have tried to mutate the codons around the early Y35 site in the WT GFP gene (i.e. around tyrosine 35) to a slower synonymous codon. As an example; tyrosine 35 was mutated from TAC (group A) to TAT (group B). When tested *in-*silico, we predicted that at least four slow translating codon mutations (two AB mutations and two AC mutations) should be introduced in order to increase protein yields (Fig. 2C, purple bars). We tested our predictions experimentally and only when four slow translating codon mutations were introduced, the WT GFP expression was rescued and showed significant increase in expression levels (Fig. 2C, blue bars). When two or three slow codon mutations were introduced, the expression levels of the WT GFP were only basal levels in both the simulation and the experiments. These results reconfirm that initial translation rates are crucial for high yields of recombinant protein expression.

### Slower transcription and translation initiation rates rescue protein expression

The use of weaker variants of both promoter and RBS to increase the average distances between adjacent mRNAs and between translating ribosomes, respectively, should reduce the probability of DITA and increase protein expression as well. We engineered weaker variant of the P70a promoter and the UTR1 RBS by introducing point mutations into the control regions. *In-vitro* transcription and translation experiments showed that the transcription initiation rate of the weaker promoter variant, P70b, was about 20 times smaller than that of the P70a promoter (Fig. S5A). *In-vitro* tests of the weak RBS variant UTR3 found that its translation initiation rate was 10 times smaller than that of the original UTR1 (Fig. S5B). Our use of either a weaker promoter or RBS enabled us to test whether DITA is affected only by transcriptional or translational density or, as our model suggests, that both factors influence the expression density, the chances for DITA and thus, the amount of expressed protein. Intuitively, the use of weaker promoter and RBS regions is expected to result in smaller amounts of synthesized protein. However, as predicted by our hypothesis and model, the counterintuitive trend was observed, according to which the weaker the control region, the higher the protein yields. This finding is true both for the weaker promoter and RBS variant, P70b-UTR1 and P70a-UTR3, respectively (Fig. 3A, purple bars). Experimental tests of this prediction showed that the weakened variants yielded up to 20 times more protein than the strong promoter-RBS construct (Fig. 3A, blue bars), suggesting that DITA can be mitigated by increasing either RNAP levels on DNA or ribosomal spacing on mRNA. Notably, when the same experiment was performed with the Y35TAG GFP mutant it showed the opposite trend, both experimentally as well as by simulation, where a weaker control region yielded less protein (Fig. 3B). Thus, by using a simple set of mutated reporter genes and incorporation of unnatural amino acids, we showed how protein synthesis yields depend, in a counter intuitive manner, on the strengths of the regulatory elements, i.e., promoter and RBS strengths, as well as on codon usage. Figure 3C is a heat map generated by the model that exemplifies the intricate relationship between promoter initiation rate and ribosomal initiation rate and resulting protein levels, it could be seen from this heat map that there is a certain set of conditions that will afford high protein yields, even for a combination of a very low promoter initiation rate and a high ribosomal initiation rate.

**Figure 3.**
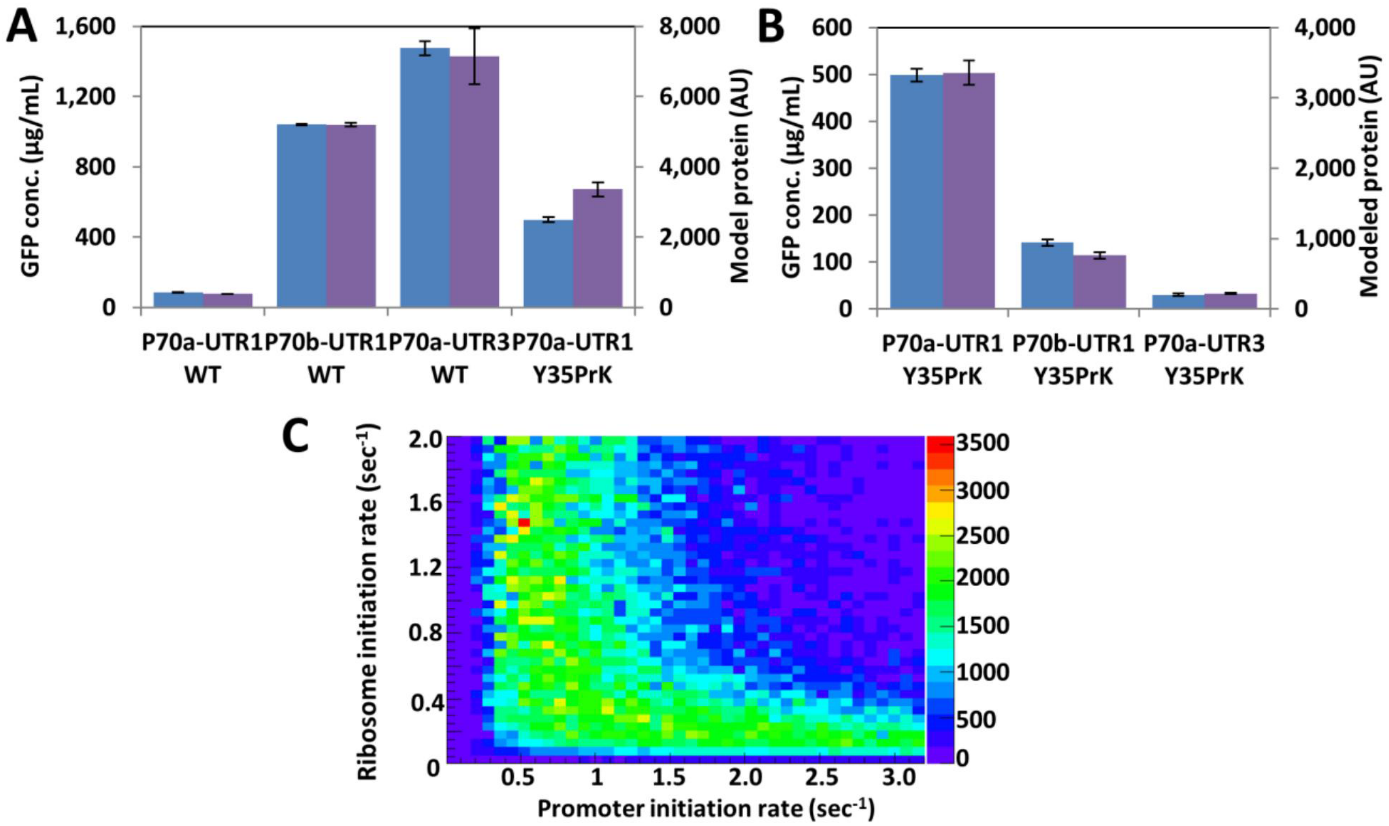
Lower initial transcription and translation rates decrease both ribosomal traffic along the mRNA as well as the chance of DITA. (A) Comparison of the experimental results (blue bars) with the modeled protein amounts (purple bars) of WT GFP expression under the control of promoter and RBS with variable initiation rates. (B) Comparison of the expression of GFP Y35PrK mutant under the control of different control regions. Experimental and the modeled results, shown in blue and purple bars, respectively. (C) Heat map of the expected amounts of WT GFP protein using different combinations of transcription and translation initiation rates.

### An analysis of mutants and initiation rate variants suggests that DITA influences mRNA levels

Under the DITA assumption, we propose that the stalling of translation somewhere along an mRNA causes all upstream ribosomes to stall while all downstream ribosomes complete translation. This hypothesis also suggests that the stretch of mRNA between the DITA site and the 3' end will be more exposed to endonuclease cleavage. For that reason, we predicted that the larger the chances of DITA, the lower the mRNA levels will be, because mRNA is more exposed to endonucleases. Using the model, we determined the amount of mRNA produced by each of the mutants and compared it to the relative quantity (RQ) of GFP mRNA found in mid-log phase cultures of the same mutants using qPCR (Fig. 4A). A comparison of the qPCR and the modeled results revealed a strong correlation, suggesting that DITA affects both protein and mRNA levels by rapidly degrading not only the mRNA, but also nascent peptides. Since high mRNA levels usually correspond to high protein expression levels, it is essential to optimize protein expression for high levels of mRNA while maintaining the half-life of mRNA by avoiding DITA. This can be accomplished by exploiting the optimal regions, in terms of transcription and translation initiation rates, for maintaining a high level of GFP mRNA and by using regulatory elements that are strong enough but calibrated to prevent DITA under high expression density conditions. The heat map shown in figure 4B is a result of a simulation of different initiation rates of the promoter and ribosomes and their influence on mRNA levels. It can be seen from the map that as expected ribosomal initiation rates have a very low influence on mRNA levels, however, our experimental results as well as the model have identified a set of conditions that mRNA levels are influenced by ribosomal initiation rates. We do not exclude other explanations for the reduction in mRNA levels such as effect on transcription initiation by the density, or a codon bias effect ^
28
^ possibly mediated by a protein ^
29
^. However, if this is the case, then it is mutually inclusive to our hypothesis.

**Figure 4.**
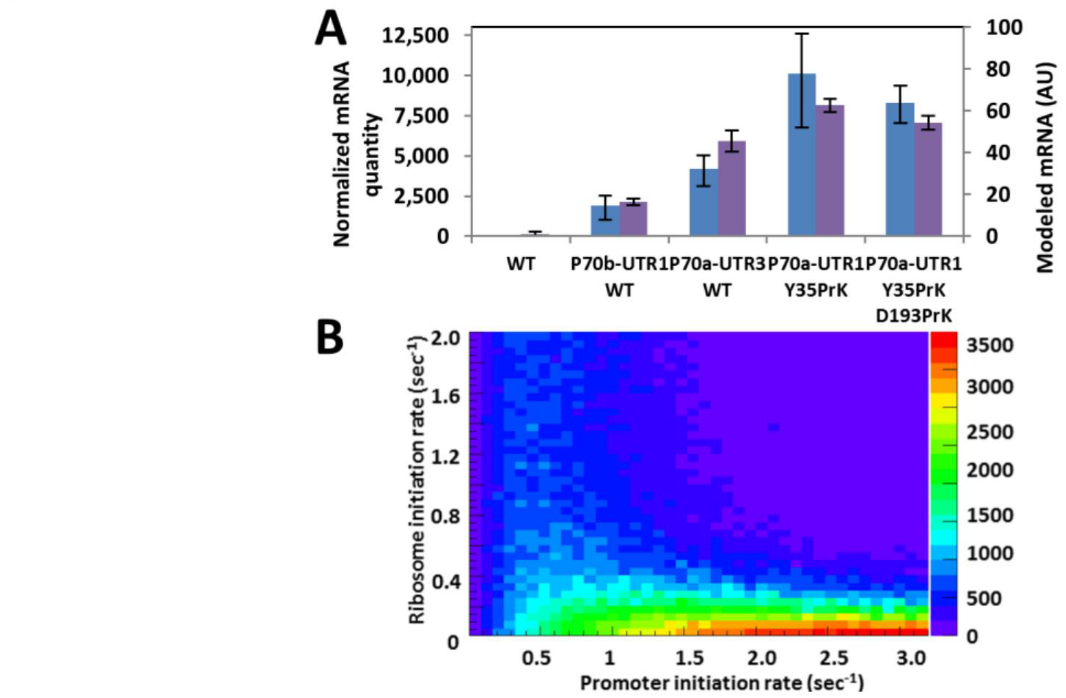
mRNA levels are also affected by DITA due to shortened half-life. (A) Comparison of the relative quantities of GFP mRNA transcripts found in mid-log phase cultures and (blue bars) with the modeled mRNA quantities (purple bars). (B) Heat map of the expected amounts of GFP mRNA transcripts using different combinations of transcription and translation initiation rates.

### Testing additional proteins supports the generality of the DITA phenomenon

To investigate whether the proposed phenomenon is a general mechanism and that it is not specific to GFP, we tested our model on three different genes: red fluorescent protein (mRFP1), *Zimomonas Mobilis* alcohol dehydrogenase II (*zmADH*) and the B1 domain of Protein L (PL), which is a small, 73-amino-acid polypeptide. The mRFP1 gene was chosen because it is a reporter protein as is GFP, however, mRFP1 shares only 26% similarity with the GFP amino acid sequence (sequences are available in the SI section), and it represents an optimized gene in terms of codon usage (it consists almost entirely of rapidly translating codons (A-type codons). The genes were tested under similar conditions to those used for GFP. The experimental results for mRFP1 were in a good agreement with the model simulations (Fig. 5A). This protein has shown the same trends as GFP both in the model and in the experiments. In contrast to mRFP1, *zmADH* is a larger, more complex gene with lower translation rates owing to its abundance of codons from groups B and C, which attenuate the translation process and result in more complex folding dynamics. The results were once more in a good agreement with the model (Fig. 5B), but we observed, contrary to model predictions, a partial rescue effect when testing expression levels with a late mutation (H297TAG). Still, a weaker promoter has given higher expression for this enzyme as well, which was predicted both by the model and by experiment. This observation is evidence that our model does not account for all factors that influence transcription/translation. Moreover, this finding suggests that co-translational folding and chaperons may introduce bifurcation points at which nascent polypeptide length is significantly reduced. Thus, the special case of a late mutation can rescue a protein from DITA. We did not address co-translational folding aspect in the current research. Lastly, PL was chosen to test the model prediction that a protein with a short polypeptide chain should have a much lower propensity for a DITA event (Fig. S3A-D). Indeed, because it is a small protein, WT PL is efficiently produced at significantly greater levels than the TAG mutated variant (Fig. 5C). The results with PL are additional experimental evidence that if the polypeptide is short enough, inter-mRNA polypeptide collision are less likely to occur although density is very high. The good agreement found between our model simulation and the experimental results for other proteins suggest that our proposed model is applicable not only to GFP.

**Figure 5.**
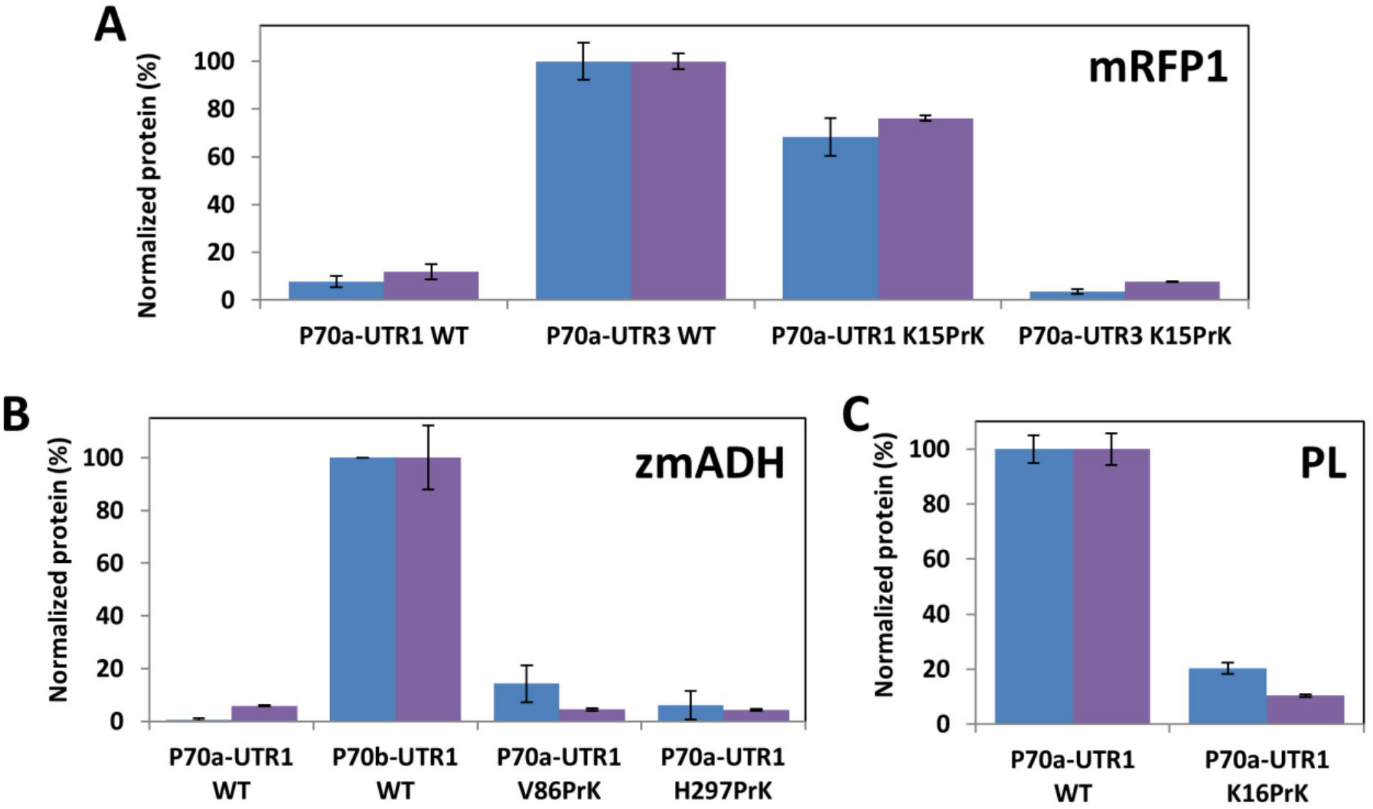
DITA is not limited to GFP and can be seen in other genes expressed using the P70a-UTR1 system and its variants. (A) Experimental (blue bars) and modeled (purple bars) protein levels ofa codon optimized WT and K15PrK mRFP1. (B) Experimental (blue bars) and modeled (purple bars) normalized expression levels of zmADH. (C) Experimental (blue bars) and modeled (purple bars) normalized expression levels of WT and K16PrK protein L (PL).

**Figure 6.**
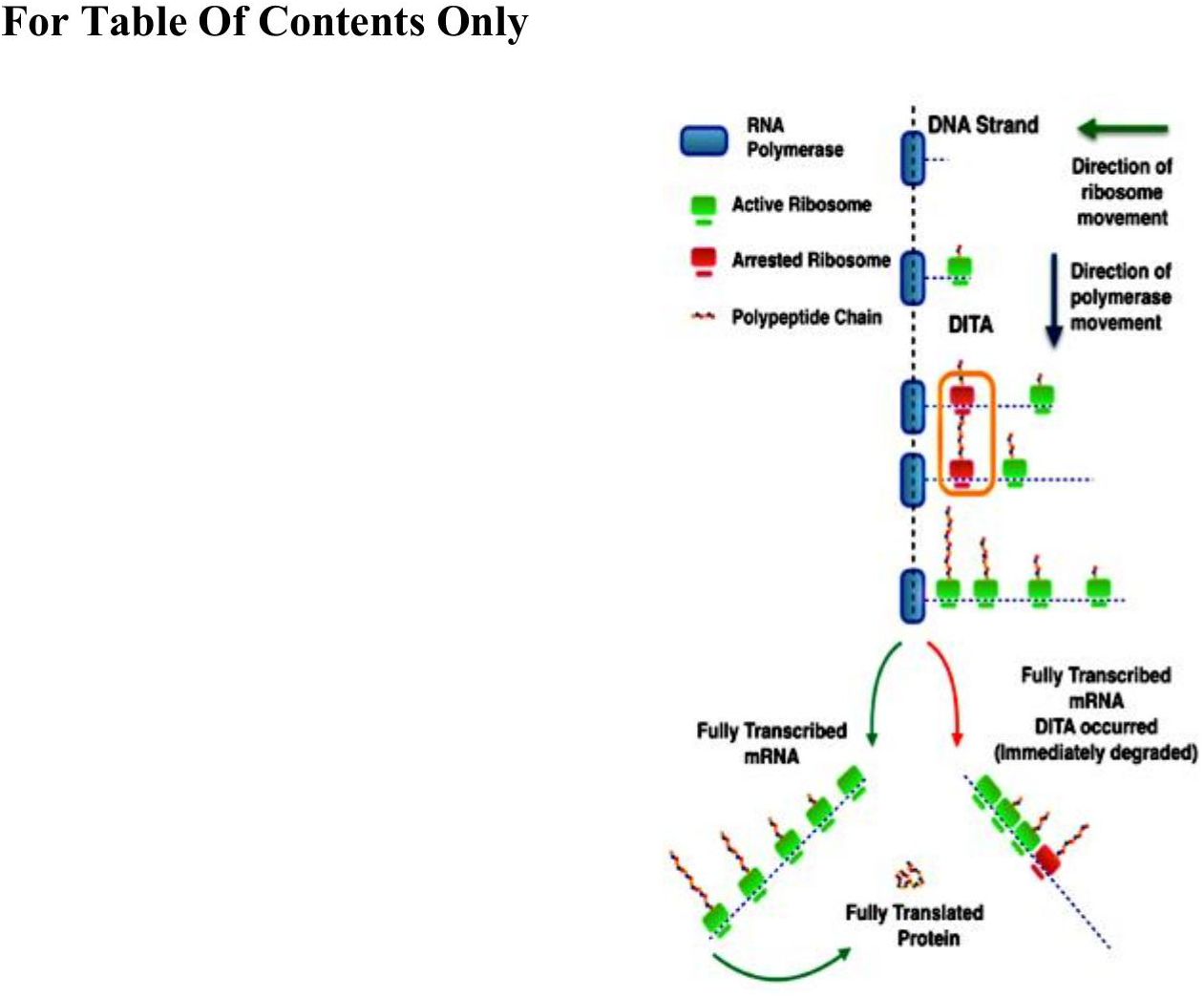

**Figure 7.**
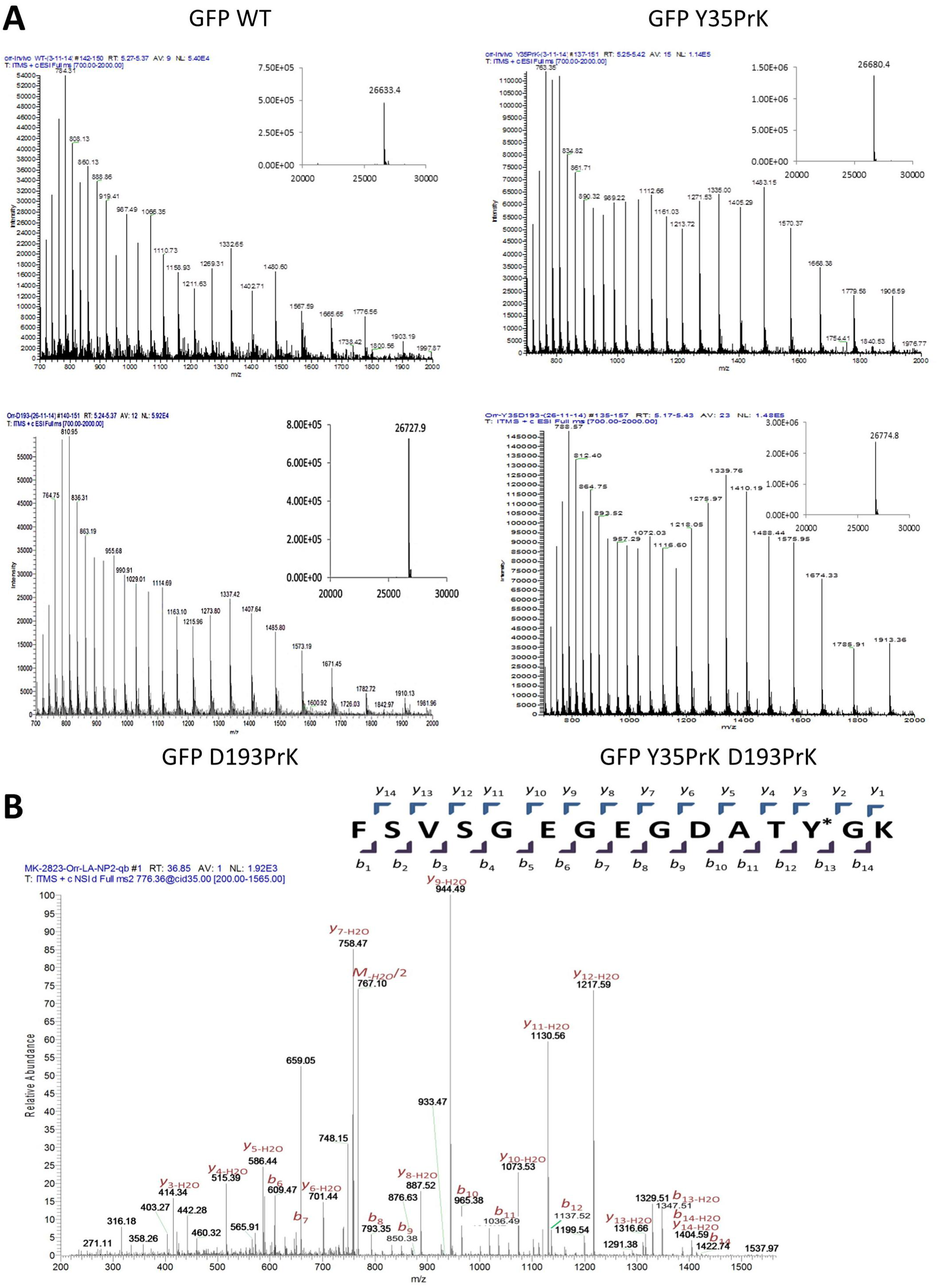

**Figure 8.**
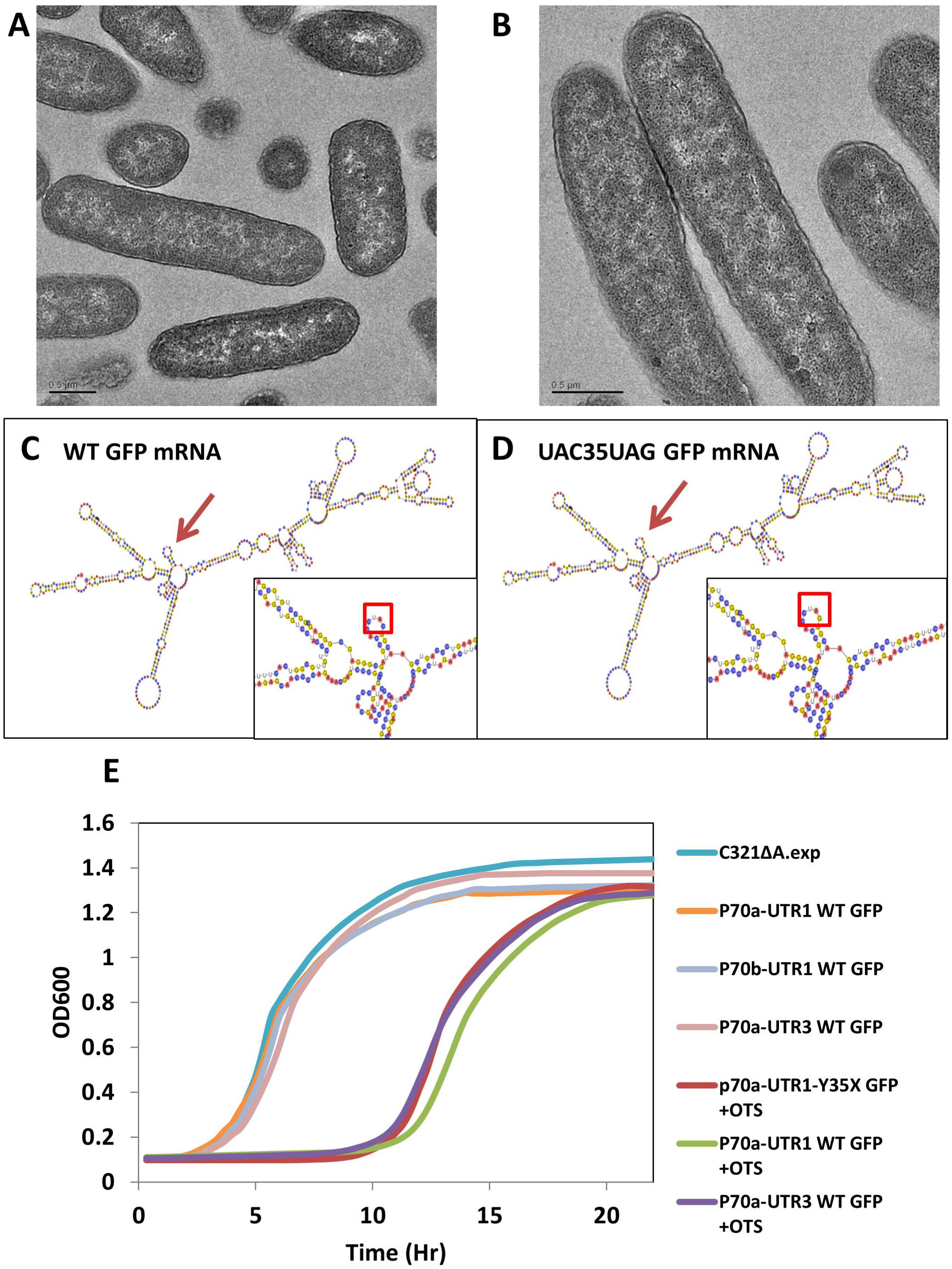

**Figure 9.**
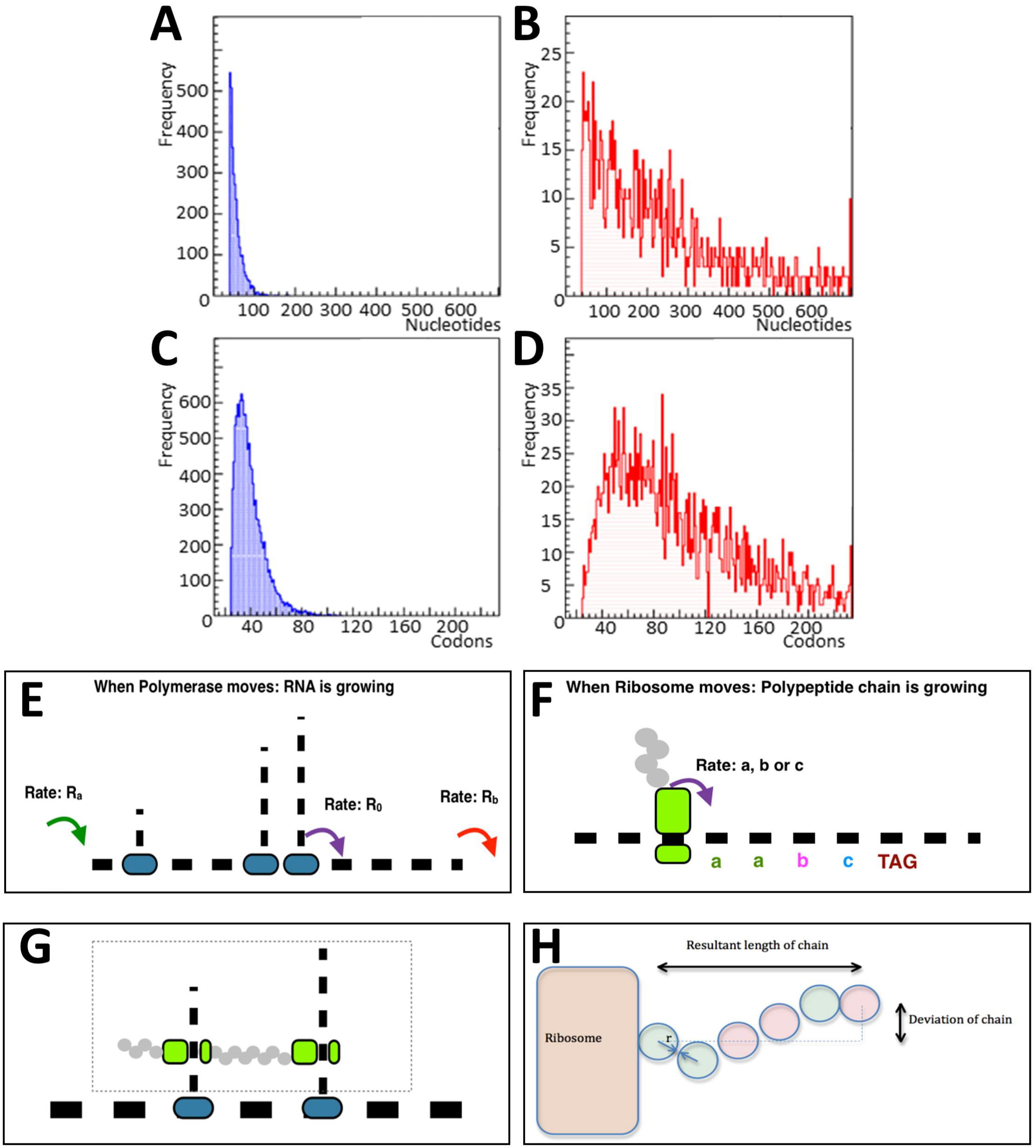

**Figure 10.**
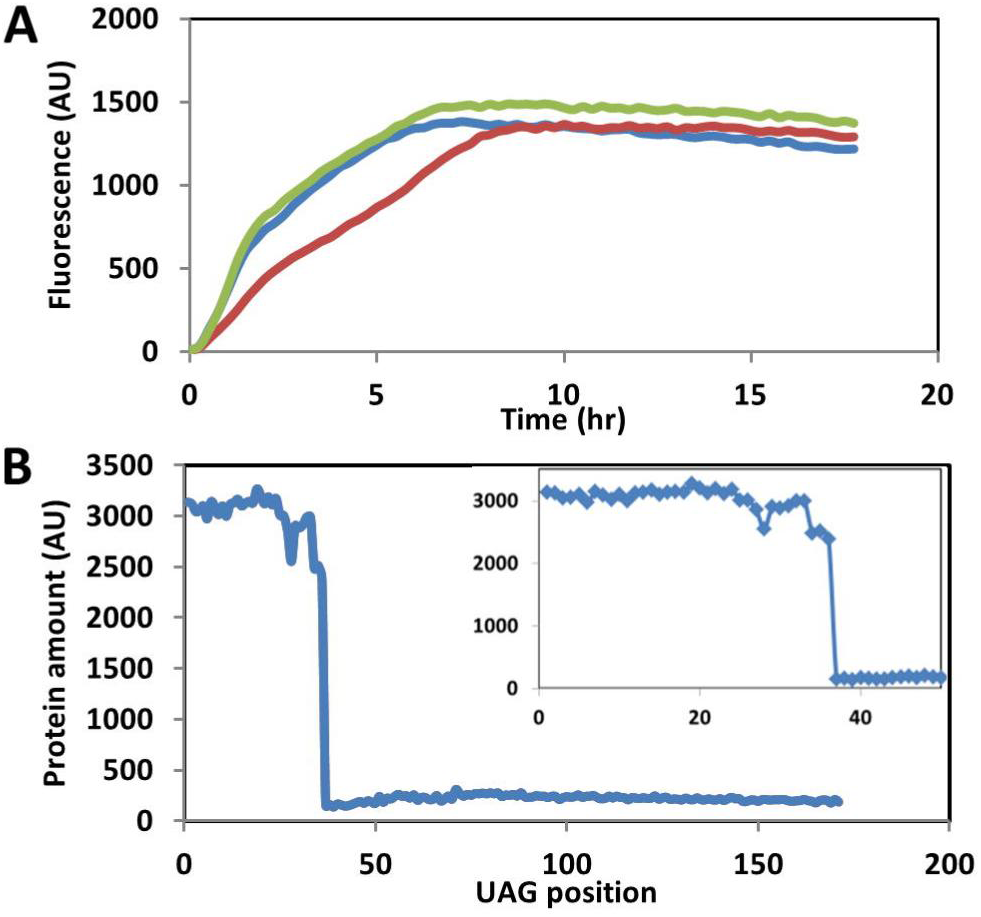

**Figure 11.**
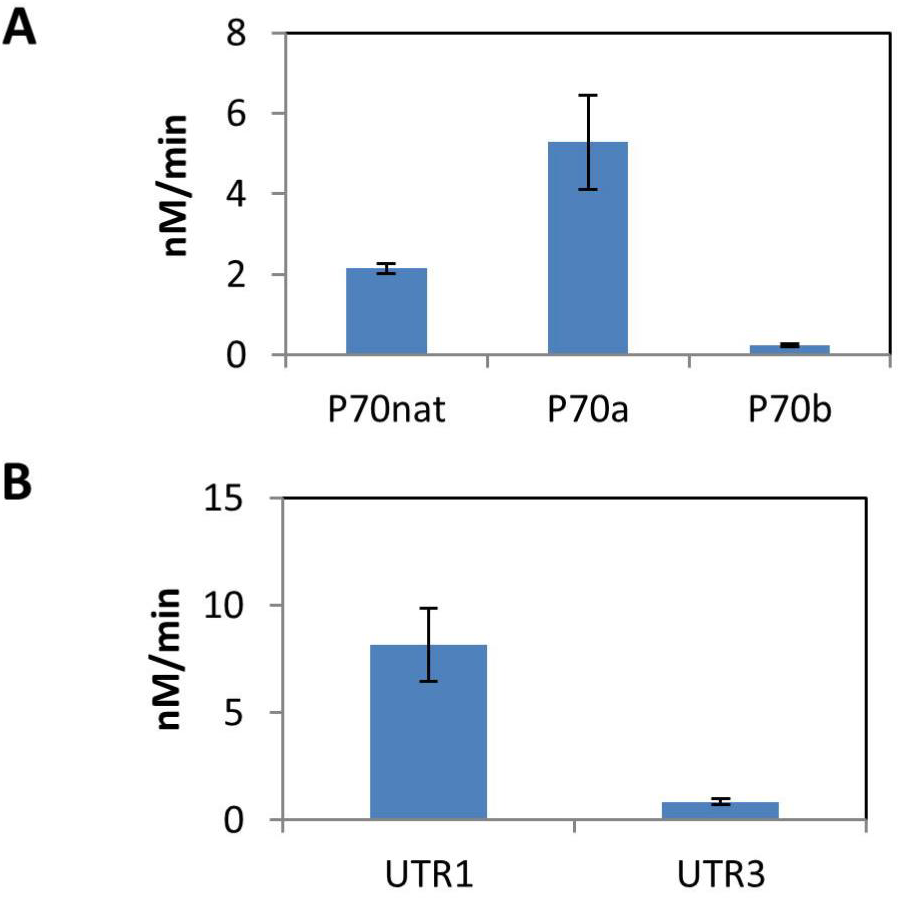

**Figure 12.**
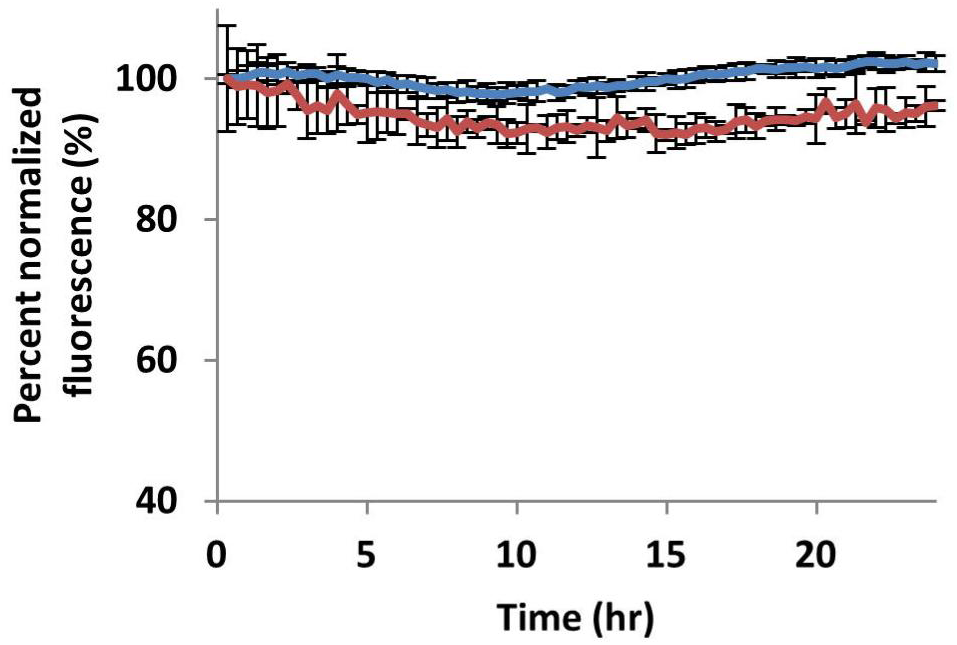

## Discussion

Due to the fact that we have no direct evidence to the occurrence of DITA, we wanted to test our hypothesis by exploring alternative explanations for this phenomenon. Alternative explanations that were excluded by us are: differences in protein stability between a protein with an ncAA and WT protein, differences in plasmid copy numbers, mRNA secondary structure differences, as well as ncAAs interference with fluorescence of the reporting protein, GFP. In order to demonstrate no apparent change in protein stability between WT GFP and Y35TAG GFP, two experiments were conducted: We monitored the stability of the WT and the mutant protein in a crude cell lysate over the course of 24 hours, showing that both proteins were stable with no significant change in fluorescence (Fig S6). In the second experiment we used synonymous, slow translating codons that were consecutively mutated around position 35, demonstrating that after the addition of four and above slowly translating codons, protein yields improve significantly to yields that are even higher than that of the protein with incorporated ncAA (Fig. 2C), these important results indicate that the same protein with no structural change, but a change in the coding sequence, can be expressed with higher yields when the initial rate of translation slows down significantly, these results are in agreement with a recent report of Zhong and coworkers^
30
^. These results also show that even with a strong promoter as is being used in this study, no hindrance from plasmid replication is observed. Evidence that attests to the fact that there is no hindrance for plasmid replication due to the existence of a strong promoter are the results with PL, since this protein is very short (ca. 70 AA) it is not affected by DITA, and high yields of expression are observed for this protein even with the strong promoter (Fig. 5C). mRNA secondary structure, could have accounted for the apparent differences in expression profiles between WT GFP and Y35GFP, however, an analysis of the mRNA secondary structure according to an algorithm written by Mathews and coworkers ^31^ have shown no difference in mRNA secondary structure. The algorithm calculates mRNA secondary structure by taking into account base pairing, free energy minimization and other thermodynamic considerations. The analysis has shown that the single nucleotide change of C→G (Fig. S2 C and D) has no implications on mRNA’s secondary structure, hence could not explain the discrepancy in expression levels. Moreover, once ribosomes bind mRNA during translation, the secondary structure is rendered almost linear, hence the predicted secondary structure is not relevant any longer and could not account for the observed difference. In order to exclude the possibility that ncAAs may interfere in any way with GFP fluorescence, we have quantified WT and mutant GFP and thus report their quantities rather than their fluorescence.

Additional possibilities were tested as well: ribosome abortion due to ribosome collisions was not excluded it could be an additional hindrance in the system but not an exclusive explanation since we could see elevated expressions of WT GFP also with a weak promoter and a strong RBS (Fig. 3A). Another possibility is that due to the strong promoter and RBS there will be an extreme consumption of translation factors (i.e. ribosomes, tRNAs, elongation factors, release factors), this possibility was excluded since it should have been seen for the much slower mutant as well (Y35PrK GFP), with the same strong promoter, multiple mRNAs will require multiple ribosomes too. Lastly, we have considered the plasmid copy number as a possible cause of low protein expression levels as is common with very strong promoters, however, our observations point to very low effect of plasmid copy numbers if any: the fact that the relatively small protein WT PL have shown high yields compared to the mutant protein using the same expression vector as for WT GFP expression, while the WT GFP have shown very small expression levels under the same conditions, contradicts the effect of plasmid copy number as the cause for low protein yields. In addition, for the same plasmid Y35PrK GFP have exhibited very high yields as well, again contradicting the effect of high plasmid copy number. Moreover, the synonymous mutations experiment (shown in Fig. 2C), demonstrates very well that after the insertion of four synonymous “slow” translating codons in the beginning of the gene, protein expression levels are recovered, for the same plasmid, yet again demonstrating that plasmid copy number could not be the cause for low protein yields.

## Conclusions

The ability of the model to accurately predict the expression trends of various proteins under different conditions led us to suggest that spatial expression density and DITA have significant effects on protein expression in cells. We note that our model does not take into account co-translational folding and therefore should not be applied to these cases. We would like to stress out that a natural system could not have been evolved to have so many strong elements to drive higher protein expression, maybe due to DITA, hence, natural systems have evolved to prevent inefficiency and energy loss. We have used artificial transcription and translation elements as well as a recombinant GFP with a synthetic sequence to demonstrate DITA. These elements were then modified to control DITA levels. In our model the expression density of any gene relies on a combination of four key determinants: translation initiation & termination rates, transcription initiation & termination rates, gene length and codon usage. Herein, we propose an additional hypothesis for the important roles of codon bias and genetic code redundancy. Although this effect was only observed in this study due to the use of highly efficient transcription and translation control regions, we infer that its effects could have significant, yet not always easy to observe implications, on the expression of all recombinant heterologous proteins. We propose that what is widely known as exogenous expression toxicity due to resource and energy depletion in some cases could be explained by DITA. In addition, we were able to show that by reducing the strength of the regulatory elements, we could lower expression density, resulting in a counterintuitive outcome that significantly improved protein yields. These protein expression dependencies were also observed at the mRNA levels of the various mutants, showing that it affects both cellular protein and *mRNA* levels, thus affecting the final quantities of protein produced. We showed that DITA occurs for several, highly dissimilar proteins, suggesting that it could be a general mechanism found in all bacteria. Moreover, our findings may also point out the importance of separating transcription and translation processes to increase the production rate of proteins, especially with longer and more complex genes. Obtaining a deep understanding of the transcription and translation processes is of an utmost importance; our findings are a novel step towards the ability to control and modify these processes, which may have a significant impact on protein expression both for fundamental research as well as for biotechnological applications.

## Experimental procedures

### GFP and mRFP1 quantification and purity assessment

GFP and mRFP1 fluorescence were measured during overnight incubation at 37°C. ncAA mutants were supplemented with PrK in a final concentration of 2 mM of ncAA. The various mutants were grown in 96 well plates while OD600 and fluorescence were measured every 20 min for up to 20 h. GFP and mRFP1 fluorescence were measured with the respective excitation/emission wavelengths of 488/510 nm and 584/607 nm. GFP mutants were purified using nickel affinity chromatography, and the resulting samples were measured using a commercial Bradford assay (Thermo Scientific, Waltham, MA). Western blot analysis was used to verify the integrity of fluorescence as a measure of protein quantity when comparing the various mutants and to eliminate the possibility of fluorescence reduction due to ncAA incorporation. For western blot analysis, goat anti-GFP and donkey anti-goat (HRP-conjugated) antibodies were used as primary and secondary antibodies (Santa Cruz, CA, USA), respectively.

### GFP purification and mass spectrometry

For the LC-MS validation of PrK incorporation, nickel affinity chromatography purification (IMAC) of 6xhis-tagged GFP was performed. Overnight cultures of 100 mL were lysed using BugBuster protein extraction reagent (Novagen, WI, USA) and 6xHis tagged GFP was purified from the crude lysate using His-Bind^®^ nickel affinity chromatography resin (Novagen). The protein-containing eluted fraction was concentrated using a Vivaspin 10 kDa cutoff concentrator (Sartorius, Göttingen, Germany) to a final concentration of 2mg/ml. The resulting concentrated fraction was analyzed by LC-MS (Finnigan Surveyor Autosampler Plus/LCQ Fleet (Thermo Scientific, Waltham, MA), using Chromolith monolithic column (EMD Millipore). The results were analyzed using Xcallibur (Thermo) and Promass (Novatia) software. MS/MS analysis was performed using standard protocols for in-gel tryptsin digestion and desalting using ZipTip µC18 (EMD Millipore). The desalted peptides were analyzed on an LTQ/Orbitrap mass spectrometer. Collision induced dissociation (CID) was used to analyze ions of interest for tandem mass spectrometry.

### *Zimomonas Mobilis* Alcohol Dehydrogenase (zmADH) expression and quantification

Cultures of C321.ΔA.exp harboring the pBEST-zmADH plasmid with the various mutants were incubated at 37°C. Cultures intended for ncAA incorporation were also supplemented with PrK at a final concentration of 2mM. zmADH expression was analyzed by quantifying ADH activity in the samples ^
32
^. The results were also semi-quantitatively verified by densitometry analysis of a western blot of the different mutants. Blotting was done using anti His-tag antibodies made in mice (Santa Cruz, CA, USA). The western blot results were analyzed using imageJ software^
33
^.

### B1 domain of protein L (PL) expression and quantification

The PL gene was subcloned to the pBEST P70a-UTR1 vector. The K16TAG mutant was created using site-directed mutagenesis (primer sequences can be found in the SI section). The two variants were transformed separately into C321Δ*Prf1.* EXP already harboring the pEVOL-Pyl OTS plasmid. The cultures were incubated overnight at 37°C in LB media supplemented with 2 mM of propargyl-l-lysine. The OD of the cultures was calibrated and lysis was performed using the protocol supplied with the BugBuster Reagent (Novagen, WI, USA). A sample of each lysate was loaded onto SDS-PAGE (WT sample was diluted by a factor of 10) and then blotted using anti His-Tag antibodies produced in mice (Santa Cruz, CA, USA). The western blot results were analyzed using imageJ software^
33
^ and the conversion to molar concentration was done using a calibration curve.

### mRNA quantification

The GeneJET RNA purification kit (Thermo Scientific, Waltham, MA, USA) was used to extract total RNA from bacterial cultures during mid-exponential phase. cDNA samples were synthesized from RNA samples using iScript cDNA synthesis kit (Biorad, Hercules, CA, USA). qPCR was performed using KAPA SYBR^®^ FAST qPCR Kit (KapaBiosystems, Wilmington, MA, USA) with the recommended relative calibration curve protocol, in the StepOnePlus Real-Time PCR System (Thermo Scientific, Waltham, MA, USA).

### Modeling the Expression density under the DITA assumption

The system was modeled in a 2D temporal-spatial model and was simulated using Gillespie algorithm. All the parameters used in the model are detailed in supplemental table S2. The parameters were assessed and determined from literature and from experimentation as described in detail in the supplemental experimental procedures section. The computational simulation enabled the assessment of mRNA and protein production kinetics and statistical assessment of the propensity for density induced translation arrest under different parameter regimes. Detailed explanation about the construction of the model including literature sources, and experiments along with the computational simulation code are available in the SI section.

### Supporting information

Supporting Information includes additional experimental procedures, 2 files with complete code of the model, gene sequences and rate maps used in the model.

## Acknowledgements

We would like to thank the following for their valuable contributions: Prof. Gilad Haran for providing us the PL gene, Prof. Yuval Shoham for providing the zmADH gene, Prof. Edward Lemke for providing us with pEVOL-PylRS plasmid, Dr. Namiko Mitarai for fruitful discussions and Prof. Kim Sneppen for invaluable advice. ERC–SG number 260647 (L. A.), The Danish Council for Independent Research (M. H. J. and M. H.) and Office of Naval Research award number N00014-13-1-0074 (V. N.) are gratefully acknowledged.

## References

(1) Browning, D. F., and Busby, S. J. (2004) The regulation of bacterial transcription initiation. Nat. Rev. Microbiol. 2, 57–65.

(2) Proshkin, S., Rahmouni, A. R., Mironov, A., and Nudler, E. (2010) Cooperation between translating ribosomes and RNA polymerase in transcription elongation. Science 328, 504–8.

(3) Chen, H.-Z., and Zubay, G. (1983) Prokaryotic coupled transcription — translation, in Methods in Enzymology, pp 674–690. Elsevier.

(4) Ellis, R. J. (2001) Macromolecular crowding: obvious but underappreciated. Trends Biochem. Sci. 26, 597–604.

(5) Tan, C., Saurabh, S., Bruchez, M. P., Schwartz, R., and Leduc, P. (2013) Molecular crowding shapes gene expression in synthetic cellular nanosystems. Nat. Nanotechnol. 8, 602–8.

(6) Mitarai, N., Sneppen, K., and Pedersen, S. (2008) Ribosome Collisions and Translation Efficiency: Optimization by Codon Usage and mRNA Destabilization. J. Mol. Biol. 382, 236–245.

(7) Jin, H., Björnsson, A., and Isaksson, L. A. (2002) Cis control of gene expression in E.coli by ribosome queuing at an inefficient translational stop signal. EMBO J. 21, 4357–67.

(8) Sørensen, M. A., Kurland, C. G., and Pedersen, S. (1989) Codon usage determines translation rate in Escherichia coli. J. Mol. Biol. 207, 365–377.

(9) Fredrick, K., and Ibba, M. (2010) How the Sequence of a Gene Can Tune Its Translation. Cell141, 227–229.

(10) Tuller, T., Carmi, A., Vestsigian, K., Navon, S., Dorfan, Y., Zaborske, J., Pan, T., Dahan, O., Furman, I., and Pilpel, Y. (2010) An evolutionarily conserved mechanism for controlling the efficiency of protein translation. Cell 141, 344–54.

(11) Cannarozzi, G., Schraudolph, N. N., Faty, M., von Rohr, P., Friberg, M. T., Roth, A. C., Gonnet, P., Gonnet, G., and Barral, Y. (2010) A role for codon order in translation dynamics. Cell 141, 355–67.

(12) Tuller, T., and Zur, H. (2015) Multiple roles of the coding sequence 5’ end in gene expression regulation. Nucleic Acids Res. 43, 13–28.

(13) Wang, L., Brock, A., Herberich, B., and Schultz, P. G. (2001) Expanding the genetic code of Escherichia coli. Science(80-.) 292, 498–500.

(14) Wang, L., and Schultz, P. G. (2001) A general approach for the generation of orthogonal tRNAs. Chem. Biol. 8, 883–890.

(15) Wang, J., Kwiatkowski, M., and Forster, A. C. (2016) Kinetics of tRNA Pyl-mediated amber suppression in Escherichia coli translation reveals unexpected limiting steps and competing reactions. Biotechnol. Bioeng. 113, 1552–1559.

(16) Fan, C., Xiong, H., Reynolds, N. M., and Söll, D. (2015) Rationally evolving tRNA Pyl for efficient incorporation of noncanonical amino acids. Nucleic Acids Res. 43, e156–e156.

(17) Lajoie, M. J., Rovner, A. J., Goodman, D. B., Aerni, H. R., Haimovich, A. D., Kuznetsov, G., Mercer, J. A., Wang, H. H., Carr, P. A., Mosberg, J. A., Rohland, N., Schultz, P. G., Jacobson, J. M., Rinehart, J., Church, G. M., and Isaacs, F. J. (2013) Genomically recoded organisms expand biological functions. Science 342, 357–60.

(18) Shin, J., and Noireaux, V. (2010) Efficient cell-free expression with the endogenous E. Coli RNA polymerase and sigma factor 70. J. Biol. Eng. 4, 8.

(19) Mankin, A. S. (2006) Nascent peptide in the “birth canal” of the ribosome. Trends Biochem. Sci. 31, 11–13.

(20) Komar, A. A. (2009) A pause for thought along the co-translational folding pathway. Trends Biochem. Sci. 34, 16–24.

(21) Woolhead, C. A., Johnson, A. E., and Bernstein, H. D. (2006) Translation arrest requires two-way communication between a nascent polypeptide and the ribosome. Mol. Cell 22, 587–98.

(22) Sunohara, T., Jojima, K., Tagami, H., Inada, T., and Aiba, H. (2004) Ribosome stalling during translation elongation induces cleavage of mRNA being translated in Escherichia coli. J. Biol. Chem. 279, 15368–75.

(23) Klumpp, S., and Hwa, T. (2008) Growth-rate-dependent partitioning of RNA polymerases in bacteria. Proc. Natl. Acad. Sci. U. S. A. 105, 20245–50.

(24) Quax, T. E. F. Claassens, N. J. Söll, D. and van der Oost, J. (2015) Codon Bias as a Means to Fine-Tune Gene Expression. Mol. Cell 59, 149–161.

(25) Mitarai, N., and Pedersen, S. (2013) Control of ribosome traffic by position-dependent choice of synonymous codons. Phys. Biol. 10, 56011.

(26) Dana, A., and Tuller, T. (2014) Mean of the Typical Decoding Rates: A New Translation Efficiency Index Based on the Analysis of Ribosome Profiling Data. G3 5, 73–80.

(27) Klumpp, S., and Hwa, T. (2008) Stochasticity and traffic jams in the transcription of ribosomal RNA: Intriguing role of termination and antitermination. Proc. Natl. Acad. Sci. 105, 18159–18164.

(28) Boël, G., Letso, R., Neely, H., Price, W. N., Wong, K.-H., Su, M., Luff, J. D., Valecha, M., Everett, J. K., Acton, T. B., Xiao, R., Montelione, G. T., Aalberts, D. P., and Hunt, J. F. (2016) Codon influence on protein expression in E. coli correlates with mRNA levels. Nature 529, 358–363.

(29) Radhakrishnan, A., Chen, Y.-H., Martin, S., Alhusaini, N., Green, R., and Coller, J. (2016) The DEAD-Box Protein Dhh1p Couples mRNA Decay and Translation by Monitoring Codon Optimality. Cell 167, 122–132.e9.

(30) Zhong, C., Wei, P., and Zhang, Y.-H. P. (2016) Enhancing functional expression of codon-optimized heterologous enzymes in Escherichia coli BL21(DE3) by selective introduction of synonymous rare codons. Biotechnol. Bioeng.

(31) Mathews, D. H., Disney, M. D., Childs, J. L., Schroeder, S. J., Zuker, M., and Turner, D. H. (2004) Incorporating chemical modification constraints into a dynamic programming algorithm for prediction of RNA secondary structure. Proc. Natl. Acad. Sci. 101, 7287–7292.

(32) Amir, L., Carnally, S. A. S. a, Rayo, J., Rosenne, S., Melamed Yerushalmi, S., Schlesinger, O., Meijler, M. M. M. M., and Alfonta, L. (2013) Surface Display of a Redox Enzyme and its Site-Specific Wiring to Gold Electrodes. J. Am. Chem. Soc. 135, 70–73.

(33) Abramoff, M. D., Magalhães, P. J., and Ram, S. J. (2004) Image processing with ImageJ. Biophotonics Int. 11, 36–42.

